# Auditory neural tracking reflects target enhancement but not distractor suppression in a psychophysically augmented continuous-speech paradigm

**DOI:** 10.1101/2022.06.18.496558

**Authors:** Martin Orf, Malte Wöstmann, Ronny Hannemann, Jonas Obleser

## Abstract

Selective attention modulates the neural tracking of speech in auditory cortical regions. It is unclear whether this attention modulation is dominated by enhanced target tracking, or suppression of distraction. To settle this long-standing debate, we here employed an augmented electroencephalography (EEG) speech-tracking paradigm with target, distractor, and neutral streams. Concurrent target speech and distractor (i.e., sometimes relevant) speech were juxtaposed with a third, never task-relevant speech stream serving as neutral baseline. Listeners had to detect short target repeats and committed more false alarms originating from the distractor than the neutral stream. Speech tracking revealed target enhancement but no distractor suppression below the neutral baseline. Speech tracking of the target (not distractor or neutral speech) explained single-trial accuracy in repeat detection. In sum, the enhanced neural representation of target speech is specific to processes of attentional gain for behaviourally relevant target speech rather than neural suppression of distraction.

## Introduction

Selective attention refers to the neural filtering processes of prioritizing relevant objects over irrelevant distractions (Desimone & Duncan, 1995). Typically, attentional selection is quantified by the difference of the behavioural or neural response to target versus distractor. However, such a difference can be driven by either target enhancement, distractor suppression, or a combination of the two. Here, we investigated how the mechanism of selective attention is implemented in neural (electroencephalographic) activity and we linked the trial-by-trial neural responses to behavioural responses associated with different sub-processes of attention.

In the visual domain, single-cell studies have shown that attention operates when multiple stimuli compete for access to neural representation. Distractors within a receptive field become suppressed, while attended stimuli are enhanced (Desimone & Duncan, 1995). The mechanism of how selective attention is implemented at the level of neural networks is still in debate in attention research (Schneider et al., 2021; van Moorselaar & Slagter, 2020). It has been argued that an often-missing, pre-defined baseline is needed to test whether the target exceeds the baseline (enhancement) and the distractor falls below the baseline (Gundlach et al., 2021; Wöstmann et al., 2022). In the visual modality, Seidl and colleagues (2012) had implemented such a “neutral” baseline by assigning a given class of stimuli as the never task-relevant, and therefore least distracting, category. They measured brain activity in fMRI (functional magnetic resonance imaging) in response to natural scene photographs that contained objects from a task-relevant (target) category, a task-irrelevant (distractor) category and a never task-relevant (neutral) category. In addition, distractor suppression was linked to attentional capture. A distractor requires to capture attention initially, followed by suppression (Alexopoulos et al., 2012; Dalton & Lavie, 2004; Gaspelin & Luck, 2018).

Speech is one of the most salient and behaviourally relevant signals in human environments, but for a long time it was not possible to study the neural processing of time-varying natural stimuli like speech quasi-continuously. Neuroscientists thus studied attention to short, isolated events due to the need for temporally discrete event-related potentials (ERP; Handy, 2005). Recently, research has begun to investigate the electrophysiology of attention to continuous speech (Ding & Simon, 2012; Lalor & Foxe, 2010; Wöstmann et al., 2017). Electrophysiological responses in cortical regions phase-lock to the temporal envelope of the speech signal (Luo & Poeppel, 2007). This linear relationship is well-captured by the so-called temporal response function (TRF), which can be interpreted as a cortical impulse response, in close analogy to the conventional ERP (Crosse et al., 2016; Fiedler et al., 2019). The TRF can indicate a stereotypical, phase-locked brain response to various acoustic features. The most often used feature is the low-frequency temporal envelope, also referred to as neural speech tracking (Obleser & Kayser, 2019). This neural speech tracking shows a robust and often-reproduced differentiation of attend versus ignored speech (Ding & Simon, 2012; Fiedler et al., 2019; Horton et al., 2013; Kerlin et al., 2010; Mesgarani & Chang, 2012). Thus, neural tracking is a feasible approach to quantify the neural processing of several speech streams at the same time to reveal the effect of attention (Ding & Simon, 2012; Puvvada & Simon, 2017; Zion Golumbic et al., 2013). In addition, Fiedler and colleagues (Fiedler et al., 2019) showed that late TRF components are associated with cortical tracking of ignored speech and are differently modulated for varying signal-to-noise ratios. These findings indicate that different components of the TRF are associated with different attentional processes. In sum, a hitherto underutilised advantage of this approach is its ability to also delineate two potential sub-processes of attention: target enhancement vs distractor suppression (Wöstmann et al., 2022).

What characterises a distractor stream in such an experimental setup? First, the implementation of the distractor stream was based on the phenomenon of negative priming, which describes the finding that a distractor from the previous trial is harder to select on the next trial (Kristjánsson & Driver, 2008; Shiffrin & Schneider, 1977; Tipper, 1985). It is assumed that a stimulus and the response it elicits become integrated into so-called “event files” in memory (Frings et al., 2014, 2020). Therefore, a specific stimulus automatically retrieves the response that was previously linked with this stimulus (Hommel, 1998). In this sense, the whole distractor stream in a given trial is distracting, since the same event that was previous task-relevant triggers a response, despite currently being task-irrelevant, and must be inhibited.

Second, it was shown that spatial statistical regularities influence selective attention on longer time scale. A location that contained a distractor with higher probability is suppressed relative to other locations. In this context, participants would learn about the location of the distractor stream and supress it over time (Wang & Theeuwes, 2018).

In the auditory modality, the attentional sub-processes target enhancement and distractor suppression have been suggested but have rarely been probed explicitly (Fiedler et al., 2019; Petersen et al., 2017; Vanthornhout et al., 2019). Here, we adopted the rationale of Seidl and colleagues (2012) and implemented three auditory speech streams, a target (task-relevant) stream, distractor stream (previously task-relevant) and – critically – a neutral stream, which is never task-relevant. Larger target-vs-neutral tracking would indicate enhancement, while smaller distractor-vs-neutral tracking would indicate suppression. Critically, it is conceivable that suppression is preceded by initial attention capture of the distractor, indicated by larger distractor-vs-neutral tracking for early neural responses (see Fig. 1B).

**Figure 1.**
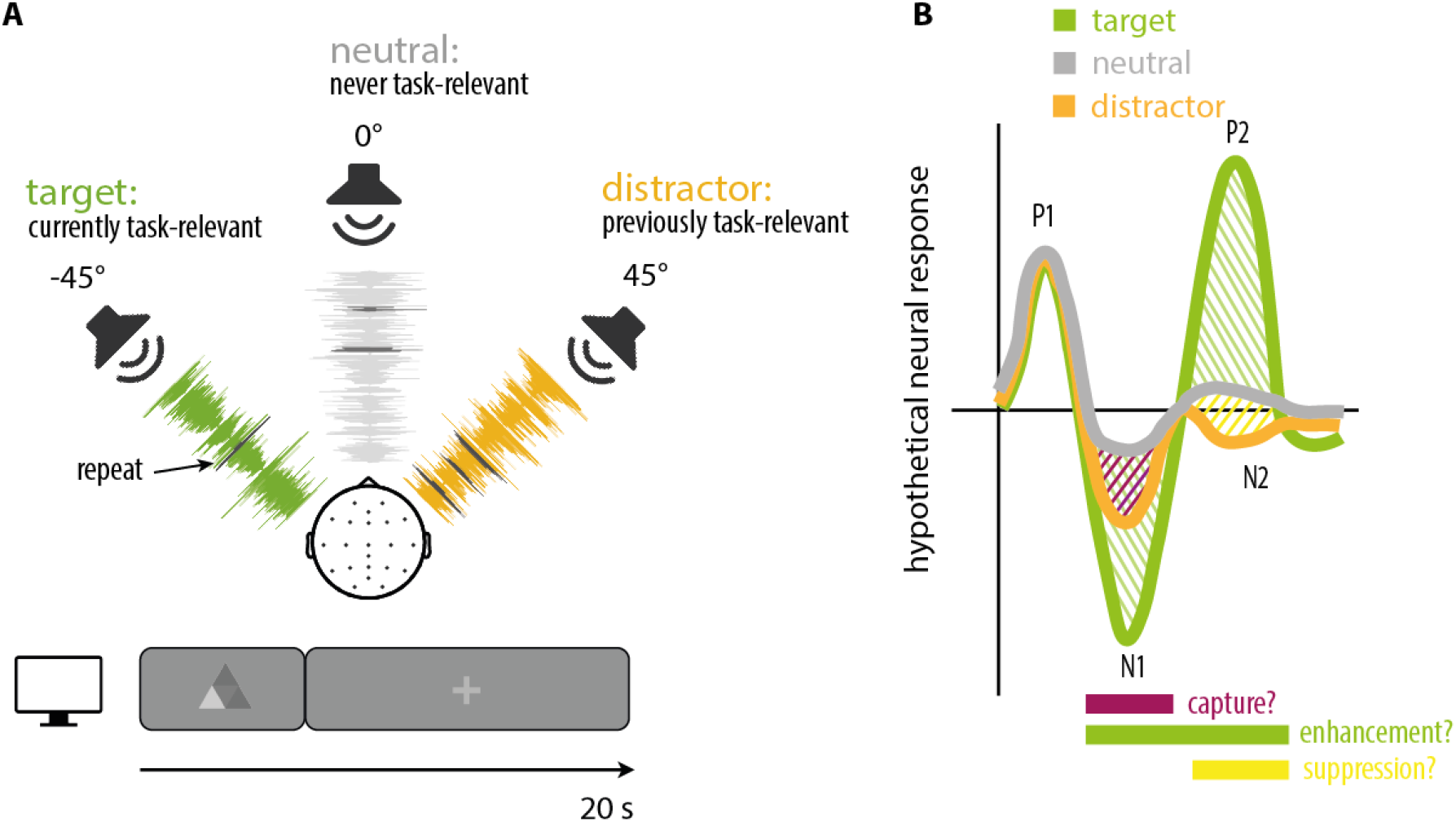
Experimental design and hypothetical results. **A**. Simultaneously, we presented three different audio streams at different locations (−45°, 0°, 45°). Participants were instructed to attend to the cued audio stream for the duration of a trial (currently task-relevant target). In the next trial, another stream was cued, which became the target stream. The stream which was previously taskrelevant became the distractor stream. During the entire experiment, the cue alternated between these two streams. The task-irrelevant (never cued) stream was defined as the neutral stream. We embedded short repeats in all three streams. Participants had to detect repeats in the target stream and had to ignore repeats in the neutral and distractor stream. Further, participants were instructed to process the content of the target audio stream. **B** Hypothetical neural outcomes. While target enhancement (stronger target-vs-neutral tracking; green) is expected for early and late TRF components, earlier components are expected to show neural capture by the distractor, that is, distraction (stronger distractor-vs-neutral tracking; red) and later components are expected to show suppression (reduced distractor-vs-neutral tracking; yellow).

However, a severe disadvantage of continuous speech paradigms thus far has been their typical lack in rich behavioural data (Hamilton & Huth, 2020). Typically, comprehension questions are asked intermittently or afterwards regarding the content of the audio stream, which are insufficient to assess task-relevance of neural responses, especially during a complex continuous speech paradigm.

In the present study, we use electroencephalography (EEG) to investigate neural responses in human participants. We asked to what extent selective attention to speech is implemented in the human brain through target enhancement versus distractor suppression, and whether enhanced tracking of target speech or suppressed tracking of distraction would explain behavioural trial-by-trial indices of attentional selection.

To this end, we designed a new experimental paradigm with two key advances over previous neural speech-tracking experiments (Fig.1A). First, a speech stimulus that was never relevant served as a neurally and behaviourally ‘neutral’ baseline, against which the processing of concurrent target speech (relevant on present trial) and distractor speech (relevant on other trials) can be contrasted (Seidl et al., 2012; Wöstmann et al., 2022). Second, listeners had the task to continuously monitor and detect short repeats in the target stream (Marinato & Baldauf, 2019) and to ignore short repeats in the distractor and neutral stream. This enabled us to contrast whether neural responses to target, neutral, or distractor speech would independently explain trial-by-trial variation in attention performance.

## Results

We recorded the electroencephalogram (EEG) from 19 young, normal-hearing participants (7 male and 12 female, mean age 21.9 years, range 18–27 years). They were presented with three continuously narrated audio streams simultaneously (Fig. 1a). On a trial-by-trial basis, they had to switch their attention between the same two audio streams. The to be attended audio stream was defined as target stream, the audio stream attended in the trial before as distractor stream and the never task-relevant audio stream as neutral stream. Participants had to detect any repetitions in the target stream as fast and accurately as possible and had to ignore the neutral and distractor stream.

Here, we analysed behavioural data in terms of signal detection theory. We tested whether selective attention is driven by an enhancement of the target, a suppression of the distractor, or a combination of the two by investigating differential neural tracking of target versus neutral speech and distractor versus neutral speech by slow (1-8 Hz) cortical responses.

### Larger repeat-evoked responses in the target stream

Overall, participants were well able to detect repetitions in the target stream (mean accuracy: 69.8% ± SEM 2.7 %; response time: 735 ms ± SEM 14.1 ms), but performance was clearly not ceiling up with up to 86 % (the highest score of single individual) correct responses (Fig. 2 a). In comparison, the false alarm rates for the neutral (mean accuracy: 2.1% ± SEM 3.4 %; response time: 789 ms ± SEM 44.7 ms) and distractor stream (mean accuracy: 2.9% ± SEM 3.4 %; response time: 801 ms ± SEM 37.7 ms) were low. Jointly, the number of hits and false alarms indicated that participants were attending to the cued target audio streams. No significant differences in response times were observed (t = 2.20; df = 30; p > 0.05, for all comparisons).

**Figure 2.**
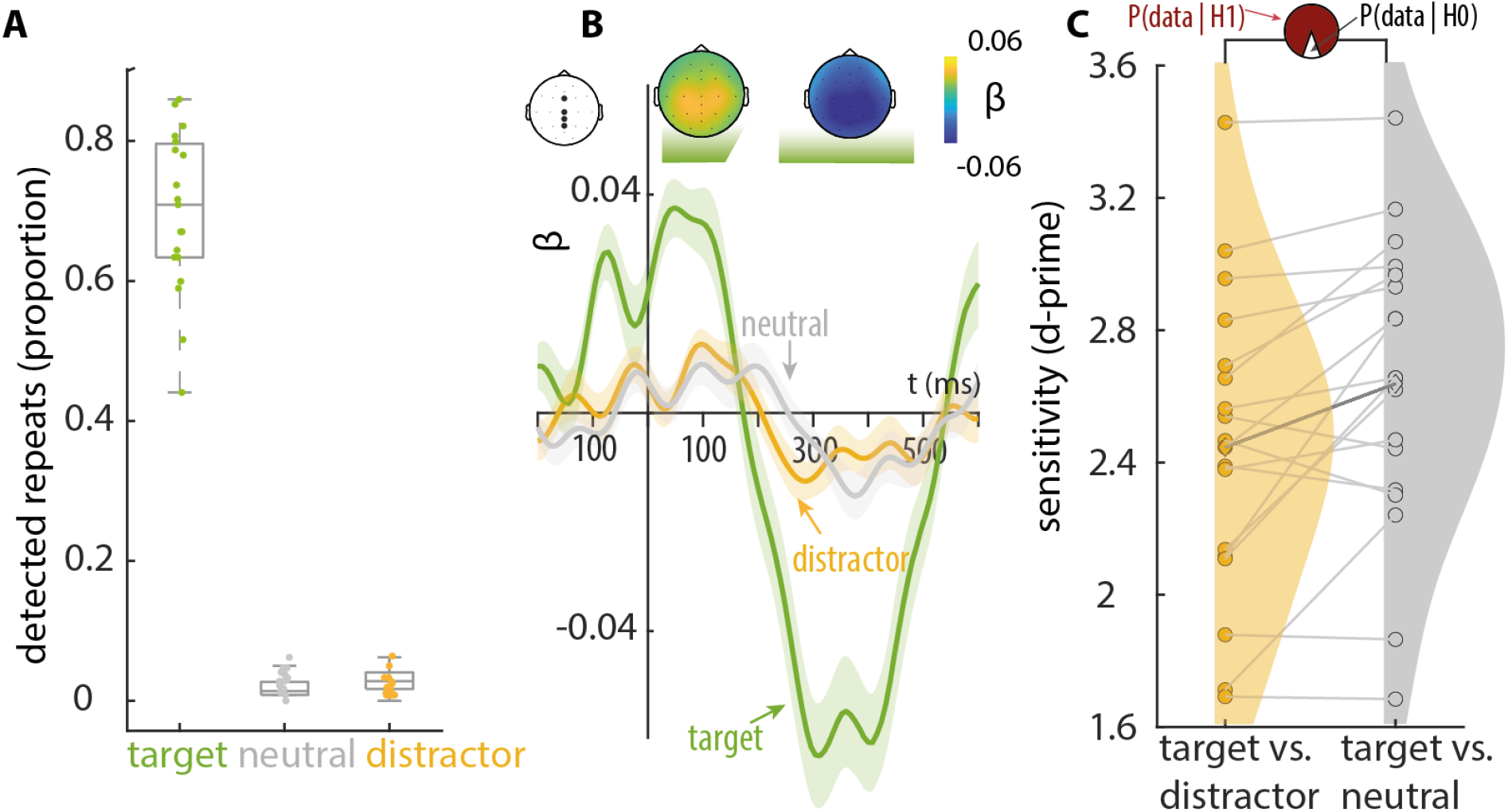
Behavioural results and ERPs to repeats. **A** Box Plots depict the proportion of detected repeats for the target (green), neutral (gray) and distractor stream (orange). Scatter dots depict individual subject data. **B** Regression based ERPs (rERPs) to repeats in the target (green), neutral (gray) and distractor stream (orange). rERPs ß-weights are averaged across subjects (N=19) and channels of interest (solid line). Shaded areas show the standard error for each time lag across subjects. Topographic maps depict ß-weights of an early time window (0-100 ms) and for a later time window (300-400 ms) for the attended stream. **C** Spaghetti plot shows the sensitivity index (d-prime) for target versus distractor stream and target versus neutral stream. Dots depict individual data and connection lines indicating data from the same subject. Shaded areas illustrate the distribution of the data. Bayes factor visualisation: probability pie charts show the ratio of the likelihood of H1(red) and H0 (white) for pairwise comparisons.

We also estimated regression-based ERPs (rERPs) phase-locked to repeat onset (Fig. 2b). rERPs to repeats in the target stream yield an auditory ERP-typical, biphasic response with an early positive deflection (0-170 ms) and a later negative deflection (170-550 ms). Topographies show ß-weights with the highest magnitude for central channels. In contrast, the rERPs for the neutral and distractor stream did not show clear rERPs. Regression based ERPs indicated a different brain response to target versus neutral repeats, but no different brain response to repeats in the neutral and distractor stream, which is in line with the observed behaviour in Fig. 2a. For further neural analysis, we treated the magnitude of these rERPs in all three streams as potential confounds and controlled for them statistically. The fact that participant’s performance was off ceiling for detected repeats in the target stream and low false alarm rate in combination with no rERPs to the distractor and neutral stream indicate that the repeats did not pop out of the streams automatically.

### Larger interference by distracting versus neutral speech

To better understand the contrast in behaviour between the neutral and distractor stream, we analysed the behavioural data in terms of signal detection theory. Based on the hit rate and false alarm rates two different d’ were calculated (Fig. 2 c). We calculated d’_target-distractor_ to index the perceptual separation of target versus distractor stream, and d’_target-neutral_ to index the perceptual separation of target versus neutral stream. Participants achieved a mean d’_target–distractor_ of 2.46 ± SEM 0.1 and a somewhat higher mean d’_target–neutral_ of 2.66 ± SEM 0.1.

A mixed model (supported by Bayesian paired samples t-test) with the regressor attention (target-distractor vs. target-neutral) statistically confirmed a significantly larger d’_target-neutral_ versus d’_target-distractor_ (t = 3.01; df = 15; p = 0.009; BF10 = 8.1, supporting H1 over H0), indicating larger interference by the distractor than neutral speech stream. A direct comparison of the false alarm rates between the distractor and neutral stream led to similar results without any transformation (t = 2.8; df = 15; p = 0.012; BF10 = 4.5, supporting H1 over H0)

### Morphology of neural responses to target, neutral and distractor speech

We analysed the neural tracking response to the target, neutral, and distractor streams by investigating the temporal, time-lagged relationship between the stimulus representation of each stream and the brain signal. This relationship is captured by an impulse response, the so-called temporal response function (TRF; see methods). Each component of the TRF is interpreted as a neural operation along the auditory pathway, analogous to the event-related potential (Davis & Johnsrude, 2003; Di Liberto et al., 2015). Here, we describe differences between the TRF for target, neutral and distractor stream, followed by statistical analysis of the neural tracking response.

As expected, the morphology of the TRF for the target stream showed the succession of P1-N1-P2 response components and the TRFs for the neutral and distractor stream showed the succession of the P1-N2 response components (Fig. 3).

**Figure 3.**
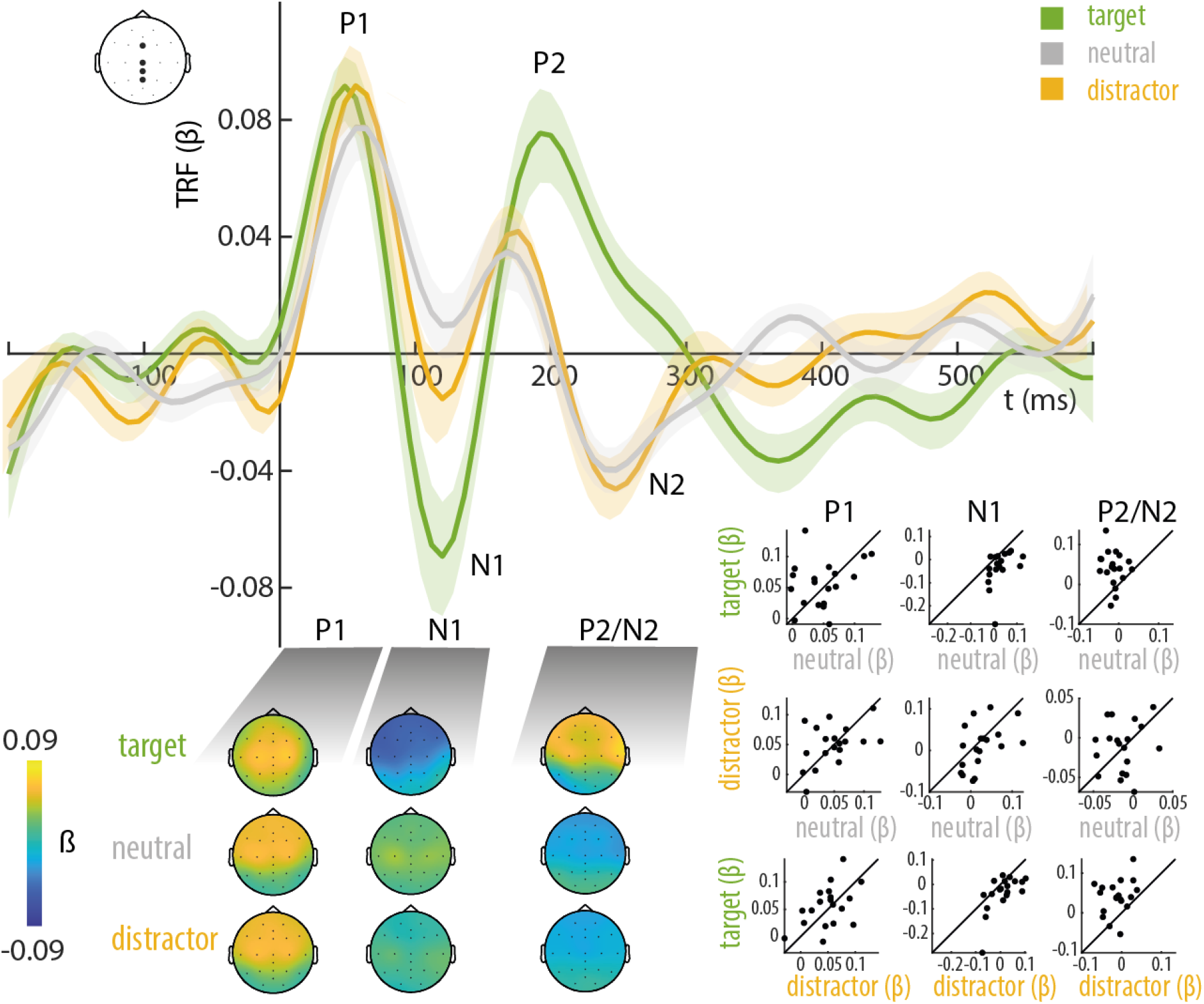
Temporal response functions (TRFs) of the target, neutral and distractor stream. TRF ß-weights are averaged across subjects (N=19) and channels of interest: Fz, Cz, CPz and Pz (solid lines). Shaded areas show the standard error for each time lag across subjects. Topographic maps depict ß-weights for time windows of the P1, N1 and P2/N2 component for the three streams. 45°-plots show the single subject (N=19) ß-weights separately for neutral versus target, neutral versus distractor and distractor versus target for the P1, N1 and P2/N2 component.

The early positive deflection P1 (0-80 ms) appeared in the TRF for the target, neutral and distractor stream without any difference, indicating no attentional modulation. Topographies (located to fronto-central regions), latencies, and polarity of the P1 component were in line with previously observed TRFs and auditory evoked potentials (AEPs) in the literature.

The later negative component N1 (80-150ms) was prominent for the TRF of the target stream. The magnitude of N1 was increased (i.e., more negative) compared with the neutral and distractor stream.

The late positive deflection P2 (170-300 ms) was only present for the TRF of the target stream. In contrast, we found a negative deflection N2 in the TRF for the distractor and neutral stream in about the same time interval. This anti-polar relationship was also reported in previous studies (Ding & Simon, 2012; Fiedler et al., 2019). However, there was no considerable difference in N2 for the TRF of the neutral stream versus the TRF of the distractor stream.

### Neural tracking reflects target enhancement, not distractor suppression

Neural tracking reflects the strength of the representation of a speech stream in the EEG (see methods for details). For neural tracking, we asked whether selective attention is driven by an enhancement of the target, a suppression of the distractor, or a combination of the two. The most important finding of this study resulted from the differential neural tracking of the target and neutral stream (target enhancement; Fig. 4b).

**Figure 4.**
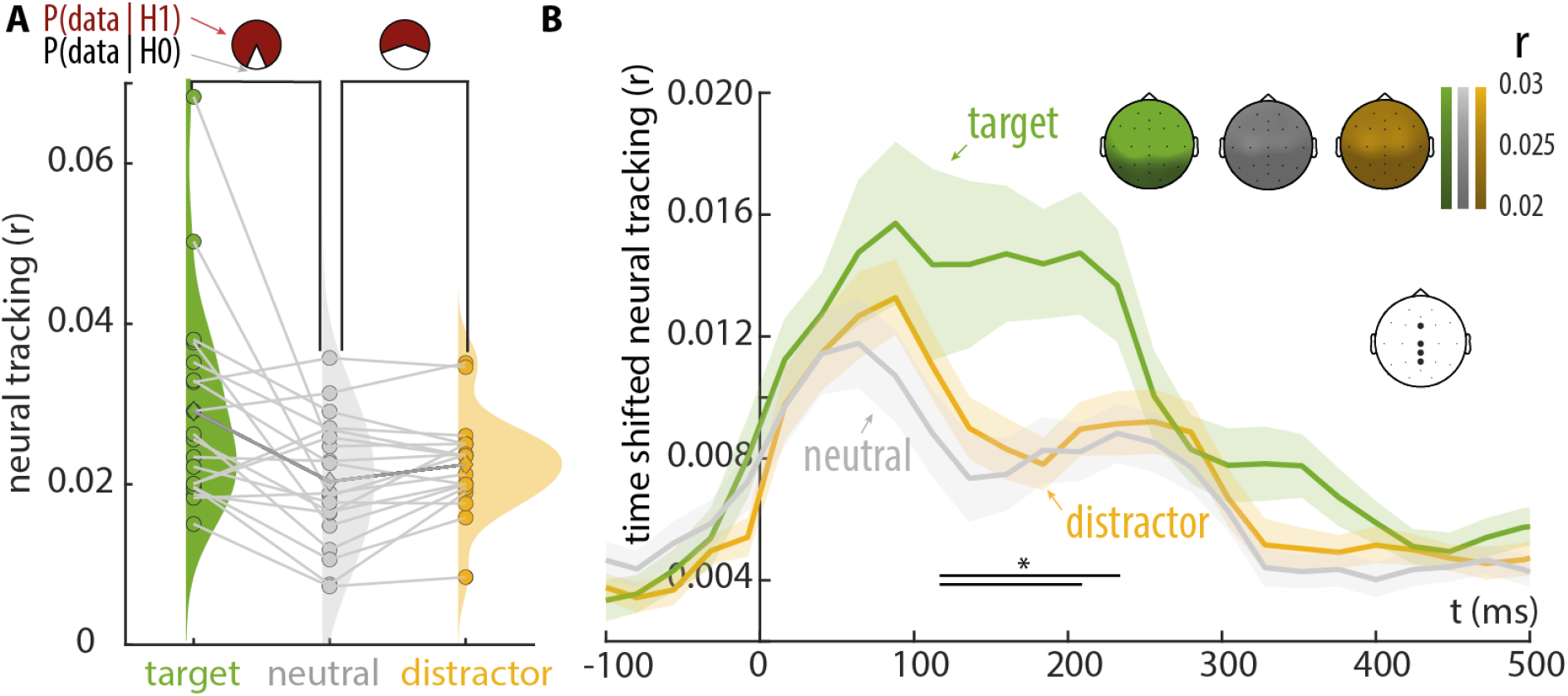
Neural tracking reveals target enhancement but no distractor suppression. **A** Neural tracking was computed based on the extracted TRFs and the envelope of the attended (green), neutral (gray) and distractor stream (orange). Spaghetti plot shows single-subject data averaged across channels of interest. Connection lines between dots indicate the same subject. Bayes factor visualisation: pie charts show probability of data given H1(red) and H0 (white) for pairwise comparisons. Shaded areas depict distributions of the data. **B** Unfolding neural tracking across time lags (−100-500 ms). Solid lines shows the averaged neural tracking (r) across subjects (N=19) and channels of interest (topographic map). Shaded areas show the standard error for each time lag across subjects. Cluster permutation test revealed two significant clusters between target and neutral (136-232 ms) and between target and distractor (136-208 ms). Black bars indicate significant clusters. No significant clusters between distractor versus neutral were found. Topographic maps depict average neural tracking (r) for the three streams (0-500 ms).

Analysis of the neural tracking (0-500 ms) revealed a difference between the target and neutral stream indicated by a linear mixed model on the mean neural tracking (0-500 ms) and Bayesian t-test for target stream versus neutral stream (t = 3.67; df = 32; p < 0.001; BF10 = 6.5, supporting H1 over H0) and between the target and distractor stream (t = 2.78; df = 32; p< 0.05; BF10 = 2, weakly supporting H1 over H0). There was no significant difference in neural tracking of the distractor versus neutral stream (t = 0.88; df = 32; p = 0.383; BF10 = 1.6, weakly supporting H1 over H0).

Unexpectedly, this effect went in the opposite direction as expected, indicating stronger tracking of the distractor versus neutral stream. Topographies revealed the strongest neural tracking for central and frontal channels.

Lastly, we analysed the temporally resolved dynamics of target enhancement and distractor suppression (Fig. 4c). Unfolding neural tracking across time lags revealed differential tracking of the target and neutral stream. Target enhancement of encoding target versus neutral speech was signified by one cluster (136-232 ms; cluster p value= 0.0044) We observed no significant clusters separating the neural response to neutral versus distractor stream.

Altogether, these findings indicate that neural tracking in a continuous speech tracking paradigm reflects a neural mechanism of target enhancement at the auditory cortical level, but no active distractor suppression.

### Neural tracking of the target stream predicts perceptual performance

To test the relationship between neural tracking and repeat detection performance, we modelled binary response behaviour (hit vs. miss) as a linear function of neural tracking for the speech streams in the target, neutral and distractor stream using a generalized linear mixed model (GLMM; see methods for details). Further, we also controlled for the different numbers of repeats in the target stream by adding trial number as a continuous predictor into the model. We also included subject ID, the number of repeats (total experiment) and the condition-to-location assignment (neutral front, left, right) as random intercepts into the GLMM.

Neural tracking of continuous speech of the target stream displayed a positive linear relationship with participant’s performance (β±SEM = 0.077±0.029; z = 2.618; p = 0.009): The higher the tracking accuracy of the target stream during a 20-s trial, the more likely participants detected repeats in that stream during that trial. We observed no such linear relationship in the neutral (β±SEM = = 0.017±0.029; z = 0.589; p = 0.556) or in the distractor stream (β±SEM = –0.023±0.029; z = –0.806; p = 0.420; Fig. 5a, left panel).

**Figure 5.**
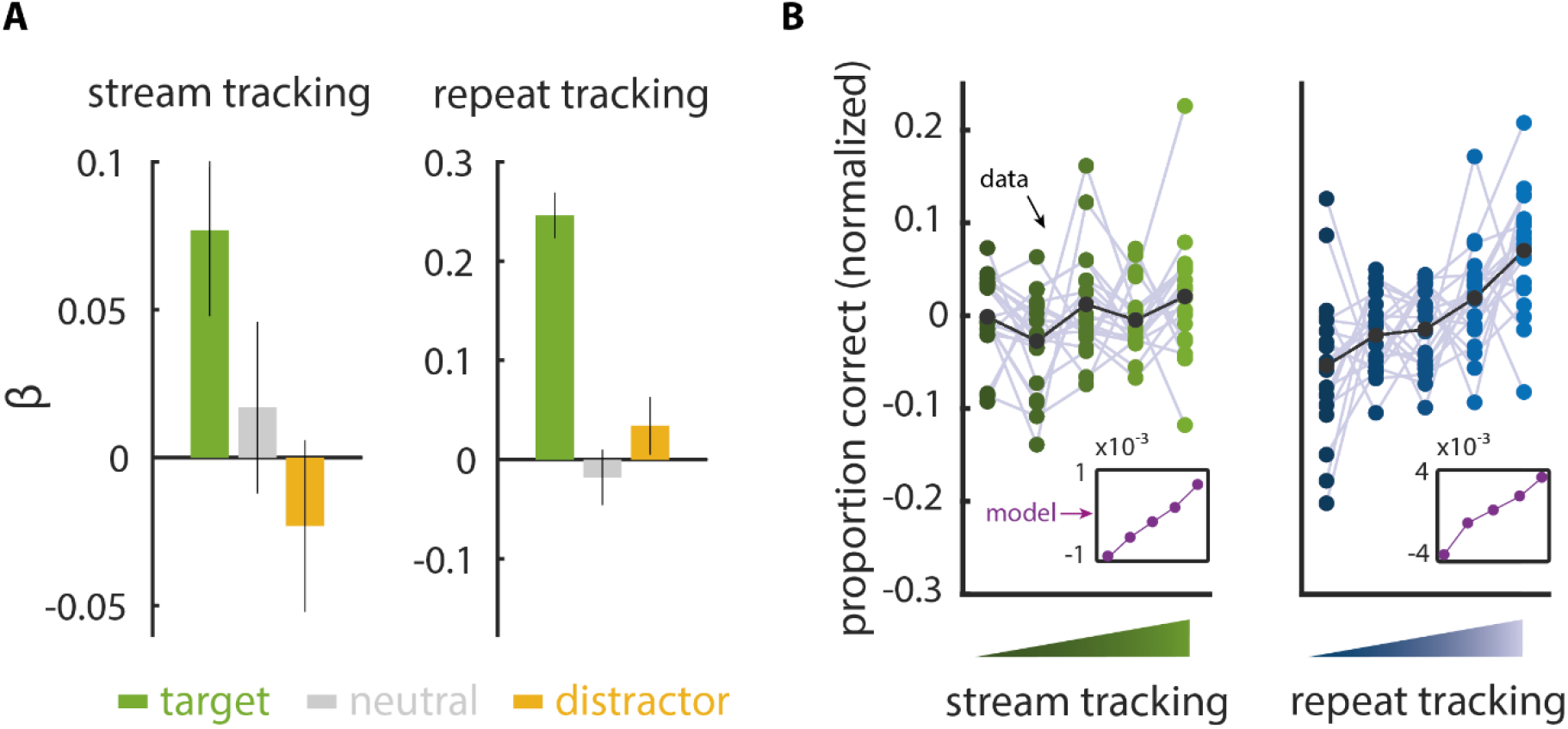
Brain–behaviour relation. **A** Standardized estimates (fixed effects, with SE) for the prediction of binary response behavior (hit vs. miss) by speech and repeat tracking for the target (green), neutral (gray) and distractor stream (orange). **B** Coloured dots and gray lines show single subject proportion correct scores; black dots and black line show the average across (N=19) subject. For illustration, data were binned by stream/repeat tracking and normalization was done by subtracting the mean of single subject data across all bins from each corresponding subject data bin. Inset shows the model prediction for each bin.

**Figure 6.**
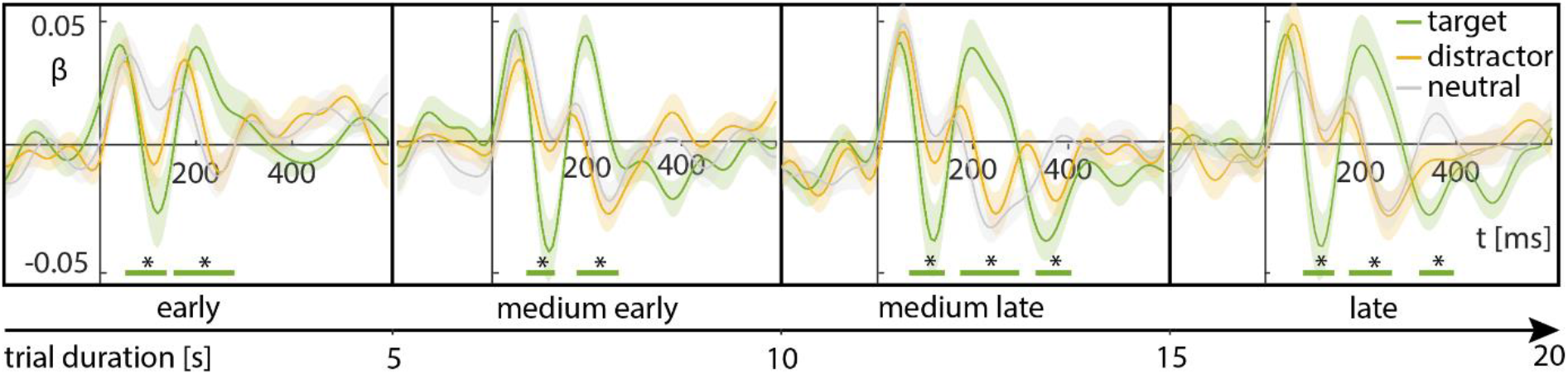
TRFs across trial duration. **A**. TRF β-weights are averaged across subjects (N=19) and channels of interest: Fz, Cz, CPz and Pz (solid lines). Shaded areas show the standard error for each time lag across subjects. TRF β-weights are estimated in four separate 5 s time windows across the trial duration (20s), representing early to late attentional processing during the trial. Cluster permutation test revealed significant clusters between target and neutral speech tracking in each time window (green bars). No significant clusters for distractor versus neutral speech tracking were observed.

Note especially that the estimates for the target stream and distractor stream pointed in opposite directions (Fig. 5a, left panel). We thus used a Wald statistic to test if the two estimates differed significantly from each other. The behaviour-beneficial contribution of the neural tracking of the target stream was positive and differed significantly from (as per sign of the estimator, behaviour-detrimental) neural tracking of the distractor stream (Z_Wald_ = 2.44, p = 0.015). As to be expected, the smaller differences of neutral versus target estimate (Z_Wald_ = –1.44, p = 0.147) and neutral versus distractor estimate (Z_Wald_ = 0.97, p = 0.332) proved not significant.

To control for potential confounding of the speech tracking in the target stream by the neural response to the to-be-attended repeats, we also included neural repeat tracking from all three streams in our model. Unsurprisingly, we observed a positive linear relationship between participant’s performance and neural repeat tracking (β = 0.246; SE = 0.023; z = 8.235; p <0.001) in the target stream. This shows that stronger neural responses to repeats in the target stream were associated with better behavioural detection of repeats. On the other hand, we observed no significant linear relationship between the tracking of a repeat in the neutral (β = −0.018; SE = 0.028; z = −0.644; p = 0.520) or in the distractor stream (β = 0.034; SE = 0.029; z = 1.197; p = 0.231; Fig. 5a, right panel). For illustration only, we binned the data by the strength of stream and repeat tracking in five bins (Fig 5b, right panel).

### Control Analysis I: Listeners process the content of competing speech streams

The behavioural outcome from the comprehension questions was not of major interest for us, since the detection of repeats provide a much more reliable and finely resolved measure of behavioural performance. However, one concern we aimed to alleviate was that participants might have been only detecting repeats rather than listening to the speech content of the target stream at all. Thus, we asked 15 multiple choice comprehension questions at the end of the experiment.

We used double iterative bootstrapping to estimate the 95% CI for the difference between the percentage of correctly answered questions and the previously determined empirical chance level of 40% (N=9 different participants only answering the questions without exposure to the full audio books; see Methods). By design, we were not able to differentiate between percentages of correctly answered questions in the target and distractor stream, as these switched their roles on a trial-by-trial basis. For instance, some question required processing on a time scale that exceed the trial length of 20s, which means that some parts of the respective audiobook content were belonging to the target and others to the distractor. Hence, we combined correctly answered questions of the target and distractor stream (50±2%, mean±SEM, range: 30-67%).

This average response accuracy was significantly better than empirical chance level (difference CI: 4.6–14.2%). The percentage of correctly answered questions of the neutral audio streams was closer to chance (48±3%, mean±SEM, range: 27–80%), but the difference to the empirical chance level was still significantly above 0 (CI: 0.9–14.6%). Percentage of correctly answered questions did not differ for the target/distractor stream versus the neutral stream (CI: –3 – 6.3%).

### Control Analysis II: Condition-to-location assignment does not confound interference by distracting speech and sub-processes of attention

In a further control analysis, we considered the possibility that the spatial condition-to-location assignment could have an indirect effect on our behavioural and neural measures. Between subjects we varied the position of the neutral sound stream (neutral: front/left/right). The different positions of the neutral stream lead also to a different assignment of the target and distractor stream. The spatial separation between the target and distractor stream was 90° when neutral was presented at 0° and 45° when neutral was presented at 45° or −45°. To control for the different spatial condition-to-location assignments, we included the factor condition-to-location assignment as covariate in our behavioural and neural analysis.

In our behavioural analysis, we observed a significant main effect on the factor condition-to-location assignment (F = 4.47; df = 15; p = 0.03). This effect is mostly driven by a significant difference between the condition-to-location assignment: neutral front versus neutral right (t = 2.96; df = 15; p = 0.01). In other words, participants correctly detected more repeats when the neutral stream was presented in the front compared with the neutral stream presented on the left or right. There was no significant difference between neutral front versus neutral left (t = 1.29; df = 15; p = 0.22) and neutral right versus neutral left (t = −1.71; df = 15; p = 0.11). Importantly, however, the difference in sensitivity was independent of the spatial position of the neutral stream: There was no significant interaction between the factors attention and condition-to-location assignment (F = 1.44; df = 15; p = 0.268).

In our neural analysis, the main effect for the factor condition-to-location assignment was not significant (F = 0.328; df = 16; p = 0.725). Importantly, the differences in neural tracking were independent of the spatial position of the neutral streams. There was no significant interaction between the factors attention and condition-to-location assignment (F = 0.88; df = 32; p = 0.482). In sum, between-subject differences in the spatial condition-to-location assignment did not confound our results.

### Control Analysis III: Unfolding of neural filters (TRFs) across trial duration

To account for the possibility that attentional processes such as enhancement, capture, and suppression unfold on different time scales over the trial duration and might cancel each other out, we divided the 20 s trial into 4, non-overlapping windows of 5s and estimated TRFs separately for each window. Cluster permutation tests revealed that target enhancement is sustained across trial duration. Importantly, we found no significant clusters for the distractor-vs.-neutral contrast (i.e., no evidence for capture or suppression). Also, a temporally more finely resolved analysis revealed no evidence for distractor capture or suppression. This analysis further supports our finding that target enhancement (i.e., attentional gain) is the dominant mechanism that modulates the neural phase-locked response to competing speech in a cocktail party scenario.

## Discussion

The present study aimed to test whether human auditory cortex enhances targets or suppresses distractors when implementing selective attention to continuous speech. To do so, we here have proposed a new, three-stream continuous-speech design with an embedded psychophysical task. The most important results can be summarised as follows:

First, the paradigm is feasible to delineate different sub-processes of auditory attention, separating a task-relevant target speech stream better from potentially neutral speech than from distracting speech. This finding proved robust under analyses controlling for stream location relative to the listener.

Second, the neural results suggest that attention is implemented through enhancement of the target stream. This lack of neural differentiation of tracking a distracting vs tracking a neutral stream speaks against mechanisms of “active” or below-baseline neural suppression of distractors at the level of human auditory cortex as measured with EEG.

Third, in line with an enhancing neural attention mechanism, the momentary neural tracking of the target but not the neural tracking of other, competing streams can predict the momentary likelihood that a listener detects events in this target stream.

### Neural tracking of speech implements enhancement, not suppression

As in previous studies (Di Liberto et al., 2015; Ding & Simon, 2012; Fiedler et al., 2019; Har-shai Yahav & Zion Golumbic, 2021; Kerlin et al., 2010; Kraus et al., 2021; Lalor & Foxe, 2010) we found the strongest neural tracking for the target stream, which was mainly due to enhanced N1 and P2 components of the cortical response. Critically extending these previous findings by implementing a neutral, task-irrelevant “baseline stream” in a three-talker paradigm, we were able to assign these previous findings to two sub-processes of selective attention: target enhancement and distractor suppression. We found a significant difference in neural tracking between target and neutral stream and no significant difference between distractor and neutral stream.

We found that participants erroneously detected more repeats in distractor versus neutral speech, which indicates attentional capture on the behavioural level. Despite this signature of capture in behaviour, we found neither suppression nor capture in the neural speech tracking response. In the visual modality, it was shown that capture and suppression go together. A distractor can capture attention, followed by suppression thereafter (Gaspelin & Luck, 2018). We have addressed this issue by analysing different time windows along the trial. However, we found no evidence for distractor capture or suppression, analysing early and late time windows separately. But that does not mean that suppression is not implemented on the cortical level in general. For instance, modulation of alpha oscillatory power is a potential neural mechanism that might implement distractor suppression in a scenario with competing auditory streams (Wöstmann, Alavash, et al., 2019).

Neural tracking of ignored speech is modulated by signal-to-noise ratio (SNR), hearing loss and perceptual demand. Fiedler and colleagues (Fiedler et al., 2019) showed that SNR manipulations of ignored speech led to differential modulation of ignored speech and the resulting neural tracking. Also, hearing loss differentially affected neural tracking of attended versus ignored speech (Petersen et al., 2017). Recently, it was found that neural tracking of distracting speech in noisy auditory scenes depends on perceptual demand (Hausfeld et al., 2021). Here following a rationale established before in visual neuroscience (Seidl et al., 2012), we manipulated the attentional fate of ignored speech by varying listener’s need to minimize or eliminate interference generated by the (previously task-relevant) distractors.

There is plenty of experimental evidence suggesting that selective attention is mainly enhancing the neural signal–to-noise ratio, thus effectively clearing or sharpening target representations in the visual and auditory domain (Desimone & Duncan, 1995; Fritz et al., 2007; Gazzaley et al., 2005; Kastner et al., 1999; McAdams & Maunsell, 1999; Mesgarani & Chang, 2012; Peelen et al., 2009; Zion Golumbic et al., 2013). In line with these findings, we show that the prioritization of the neural representation of the target auditory input is mainly implemented by an enhancement of the target. In this respect, our results are also notably in line with a recent visual EEG study on attentional suppression by Gundlach and colleagues (Gundlach et al., 2021). Also, another recent study investigated whether exogenous attention led to facilitation of attended information, suppressed unattended information, or both (Keefe et al., 2021). Both studies found that attention rather operates on target enhancement than distractor suppression.

Generally, our study adds to the unsettled debate in attention research over neural implementations of suppression. Even before the present study, evidence in the literature for distractor suppression has been mixed, with some studies speaking to (Desimone & Duncan, 1995; Schwartz & David, 2018; Seidl et al., 2012; Wöstmann, Alavash, et al., 2019) and others speaking against distractor suppression (Gundlach et al., 2021; Keefe et al., 2021; Noonan et al., 2018).

Classical theories of attention permit some form of distractor suppression (Broadbent, 2013; Treisman, 1964), and there might well be distinct types of distractor suppression as endpoints to a continuum. Also, from a neurocognitive vantage point, distractor suppression does not need to be one single process and could be rather implemented via multiple neural mechanisms.

Firstly, suppression could be driven by the current intention of the observer extracting statistical regularities of certain features such as location of a distractor over time, enabling the brain to learn to produce suppression (Wang & Theeuwes, 2018; Wöstmann et al., 2022). In the long term (duration of the experiment), participants could learn based on statistical regularities the location (same location of distractor stream) and the voice of the talker (same voice). Secondly, in the short term (every trial), participants are cued (current intention) to attend to one stream and to suppress the distractor (negative priming). In principle, our paradigm might initialise both of these types of distractor suppression. While it is debatable whether the effect of our negative priming manipulation persists over the whole trial duration (probably decreasing over time), learning and using statistical regularities of the distractor over time should persist in the long term of the experiment. However, we found no significantly suppressed neural tracking of the distractor vs. neutral stream, which suggests that the neural speech tracking response does not implement distractor suppression. Contrary to our hypothesis, results hinted rather at a potentially stronger tracking of the distractor compared to the neutral stream although this was not a statistically robust observation in the present data. For future studies, it is nevertheless important to consider such an attentional capture of the distractor stream (Gaspelin & Luck, 2019).

Secondly, distractor suppression can be generally divided into proactive (processing before the distractor appears) and reactive suppression (processing after the distractor has captured attention; Chelazzi et al., 2019; Wöstmann et al., 2022). The amplitude of neural alpha oscillations (~10 Hz) related to top-down selective attention processes can be modulated by target- and distractor-processing. Wöstmann and colleagues (Wöstmann, Alavash, et al., 2019) found that alpha power during the anticipation of competing tone sequences implements distractor suppression independent of target enhancement. Their results speak to a proactive implementation of distractor suppression. But neural tracking is characterized by the time-lagged neural responses which phase-lock to the stimulus. Due to this characteristic, neural tacking is rather suited to investigate reactive suppression than proactive suppression. With respect to these distinguishable sub-processes of distractor suppression, our results indicate that at least reactive suppression is absent for auditory cortex responses in a multi-talker situation.

### Auditory attention exploits statistical regularities to separate distracting versus neutral speech

When considering how distracting versus neutral, task-irrelevant speech might be encoded neurally, a previous auditory study using also three streams had suggested that higher-order auditory areas provide an object-based representation for the foreground, but the background remains unsegregated (Puvvada & Simon, 2017). At first glance, our results are broadly in line with this conclusion, but note that Puvvada and Simon had not applied any differential task manipulation to the two background speech streams, which we aimed to achieve here. The here proposed experimental paradigm aimed to strike important compromises in studying the listener’s neurocognitive ability to separate target, distractor and neutral speech.

In contrast to trial-based designs, continuous speech paradigms often lack rich behavioural data. Usually, comprehension questions regarding the content of the audio streams are asked to differentiate between attended and ignored audio streams (Broderick et al., 2018; Fiedler et al., 2019). Asking comprehension questions has some drawbacks. Comprehension questions usually refer to a comparable long-time range. This limits the number of questions and thus the number of behavioural data that can be extracted from the experiment. Further, in our paradigm participants had to switch their attention every 20s between two audio streams, which did not allow us to strictly assign the question to attended or ignored parts of the audio streams. Hence, it was insufficient solely to ask comprehension questions to investigate the listener’s cognitive ability to separate target, distractor, and neutral speech on the behavioural level. More fine-grained behavioural data were needed ideally without losing much of the ecological validity of natural speech.

We used short repeats in the audio streams to obtain rich behavioural data. In trial-based designs, participants are asked much more frequently to respond, which also ensures a steady engagement into the listening task. Baldauf and colleagues (2019) also embedded short repeats in auditory objects, arguing that such a detection task requires the processing of the acoustic stream at the level of auditory objects. Such a repeat detection task might thus be particularly suited to study object-based mechanisms of selective attention. Adopting this approach here, we found that participants detected much more repeats in the target (hits) compared to the neutral and ignored stream (false alarms).

Recall that, in our paradigm, participants had to switch attention between the same two streams while they had to ignore the never-task relevant neutral stream. Importantly, we found a significantly larger behavioural interference by distractor speech than by neutral speech, but what is the underlying mechanism? Our results suggest that the neural fate of a stream on the previous trial has the potency to make it more distracting and captures attention on the text trial. This corresponds with the concept of negative priming. Negative priming refers to the effect that the reaction to a stimulus that was previously ignored is more error-prone and slower (Tipper, 1985). Classical negative priming designs consist of two main components: prime (trial N) and probe (trial N+1). The prime presents a certain stimulus (or stimulus feature) as a distractor, which becomes the target in the probe trial. Negative priming has been studied in vision in a detailed manner (Fox, 1995; May et al., 1995).

Although there are fewer studies that investigated negative priming in auditory selective attention, they reported similar results (Frings et al., 2014). Nowadays most researchers agree that auditory negative priming (similar in vision) is explained by inhibition and retrieval theories (Frings et al., 2014). Longer response times and higher error rates are typically observed relative to a no priming condition (Banks et al., 1995; Mayr et al., 2011; Mayr & Buchner, 2010). Notably, we did not present the same segments of the audio streams on two consecutive trials. Participants had to attend and ignore different segments of the audio streams in each trial, due to the ongoing structure of continuous speech. We assume that it was rather the spatial location or/and the voice that was associated with negative priming and leaked into the present trial, than the identity of the auditory stimulus. On the one hand, if a listener attended to a specific feature of an auditory object, not only this specific feature is enhanced, but all features related to the selected object (for review, see Shinn-Cunningham, 2008). On the other hand, one could argue that this also holds for features concerning negative priming and object suppression.

A more recent study varied randomly the location of the target and distractor and the speaker (Eben et al., 2020). They demonstrated negative priming in auditory selective attention switching with the spoken material. In sum, our new paradigm has proven feasible to utilise the negative priming phenomenon to unravel listeners’ separation of distractor speech versus neutral speech.

### Neural tracking of target but not distractor explains performance

Continuous speech paradigms often lack rich behavioural data. But only if we unravel the precise relationship between brain and behaviour can we reach a veridical understanding of cognitive processes such as selective attention (Krakauer et al., 2017). We embedded short repeats into the speech streams which served as a trial-by-trial measure for behaviour. In addition, this also enabled us to predict behaviour from neural responses on a single-trial level. We found that neural tracking of the target stream only predicted trial-by-trial variation in repeat detection. Our results not only provide support to the functional relevance of neural speech tracking (Tune et al., 2021), but significantly expand this by providing an explanation for the underlying sub-processes of auditory selective attention, that is, enhancement of the target and not suppression of distractors predicts performance. In addition, this finding supports the feasibility of our new continuous speech paradigm since we found a significant relation between the neural tracking of continuous speech and the repeat detection behaviour. Further, the finding supports our previous findings since only target enhancement predicts behaviour. Indicating that the prominent process of selective attention is target enhancement rather than distractor suppression.

### Limitations

There are three limitations regarding the operationalization of the neutral and distractor stream. First, the attentional manipulation by their respective task-relevance (Seidl et al., 2012) of the distractor stream might not lead to an interference strong enough that distractor suppression was useful. Thus, it is possible that negative priming in combination with the spatial and/or spectral separation of the audio streams was insufficient to activate the need of distractor suppression in our study. Future studies could address this by varying for instance the separation between the audio streams (Hausfeld et al., 2021). The task may become more difficult with smaller spatial separation, which potentially activate distractor suppression.

Second, the neutral stream can be conceived as a weaker distractor. We operationalized the neutral stream as the never task-relevant stimulus. However, the neutral stream is not neutral in the strongest sense: Like the distractor stream, it was associated with the attentional background since it had to be ignored by the listener (Puvvada & Simon, 2017). In other word, the neutral stream was more similar to the distractor stream compared to the target stream.

Third, our sample size (N = 19) could have been too small to detect small distractor suppression effects. Note, however, that any such distractor-suppression effect size would need to be put in perspective to the considerable effect sizes of target enhancement we observed. So, the relative conclusion about target enhancement vs distractor suppression would remain. Thus, the conclusion stands that target enhancement is the behaviourally and neurally more prominent sub-process of selective attention in a continuous speech paradigm.

## Conclusion

In attention research, previous paradigms have rarely aimed at separating mechanisms of distractor suppression conclusively from target enhancement. Using a new, psychophysically augmented continuous-speech paradigm with three speech streams, our results demonstrate that neural tracking of continuous speech reflects target enhancement, not distractor suppression. These findings call for a refinement of our models about enhanced neural responses to speech and should account for a specific sub-process of selective attention, that is, the enhancement of targets rather than the suppression of distraction.

## Methods

### Participants

Nineteen young adults (12 female, 7 male), aged between 18 and 27 years (Ø 21.9) participated in the present study. All participants had German as their mother tongue and reported normal hearing and no histories of neurological disorders. To verify normal hearing, we measured pure tone audiometry within a range of 125 to 8000 Hz. All participants showed auditory thresholds below 20 dB for the tested frequencies. They gave written informed consent and received compensation of 10€/hour. The study was approved by the local ethics committee of the University of Lübeck.

### Stimulus materials and spatial cue

We presented three different narrated book texts as audio, spoken by different male, professional talkers (“Michael Kohlhaas” by Heinrich von Kleist, “Pole Poppenspäler” by Theodor Storm, and “Das Wrack” by Friedrich Gerstäcker). We chose audio streams that were fictional instead of fact-based, to minimise the impact of variations in prior knowledge on a topic and a resulting possible bias to one of the audio streams. All three audio streams overlapped in time, at an average intensity (mixture) of about 65 dB(A), which matches normal conversation levels.

The following processing steps of the stimuli were done using custom written code in MATLAB (Version 2018a Mathworks Inc., Natick, MA, United States). The sound files were sampled with 44.1 kHz and a 16-bit resolution. The sound level was matched to the same long-term root-mean-square (rms) dB full scale (dBFS) between the three audio streams. Silent periods were truncated to maximally last 500 ms (O’Sullivan et al., 2015).

We embedded short repeats in the audio streams by pseudo-randomly selecting a 400-ms segment from the original stream and repeating it directly thereafter (Marinato & Baldauf, 2019). The first repeat was presented at least two seconds after stimulus onset. Each repeat was included in the sound stream by a linear ramping and cross-fading. The linear ramping was done by using a window of 220 samples (5 ms) of the end of the to be repeated part (down ramp) and by using the first 220 samples (5 ms) of the repeat itself (up ramp). The cross-fading was done by adding the down and up ramp together.

The onset time of each repeat was drawn randomly to avoid predictability of the repeat: To avoid that repeats occurring in the different streams overlap in time, the distance between two repeat onsets was at least 2 seconds.

We further used a rms (root mean square)-criterion (rms of the repeat had to be at least the same rms as the stream of which the repeat was drawn from) to avoid undetectable repeats of low sound intensity.

The cue was presented at the center of the screen (resolution: 1920×1080, Portable HDMI Screen, Wimaxit) in front of the participant (distance: 1 m). The cue (Fig. 1a) consisted of three sub-triangles which had a size of 1.3° visual angle pointing to the three sound sources (front, left, right). The background of the screen (RGB: 127, 127, 127), the cued sub-triangle (RGB: 204, 204, 204) and the not cued triangles (RGB: 115, 115, 115) were kept in different shades of gray to keep the contrast low. The bright triangle indicates the to-be-attended position. Since the cue and the fixation cross were presented at the same time as the auditory stimuli, we ensured that the possible interference between visual and auditory neural responses was as small as possible. To this end, the change between the fixation cross and cue was made smooth by linearly fading in and fading out (50 ms each) the cue.

### Experimental Setup

The experiment took place in a laboratory space with eight loudspeakers (Genelec: Speaker 8020D, Denmark) arranged in a circle with a radius of one meter. The loudspeakers were spaced at 45 degrees. A chair was placed in the middle of the radial speaker array, face-aligned to the loudspeaker at position 0°. The three audio streams were presented over the three frontmost loudspeakers (−45°, 0°, 45° in the azimuth plane, elevation was not adjusted for participants’ height, ground to loudspeaker distance: 1,20 m, the five remaining speakers were not used in the present experiment). In advance, participants were briefed about the experiment. Importantly, they were not briefed about the condition-to-location assignment of the streams. Each participant was asked to keep eyes open, focus to the centre of the screen and to sit as relaxed as possible. To avoid head motion, a chin rest was used. The height of the chin rest was adjusted for each participant.

### Experimental Procedure

We created a new experimental paradigm to investigate the underlying neural mechanism of selective attention (Fig. 1). The experiment was designed using MATLAB (Version 2018a Mathworks Inc., Natick, MA, United States) and Psychophysics Toolbox extensions (Brainard, 1997; Kleiner, 2007; Pelli, 1997). Participants were presented with three concurrent audio streams. Each trial started with a cue. The cue indicated which location to attend. The cue was presented for 500 ms. After the cue, a fixation cross was presented for the remaining trial duration (19.5 s). However, the auditory stimuli were presented simultaneously with the cue and the fixation cross resulting in a continuous playback of the auditory stimuli without any breaks between trials. Hence, the next trial started instantly after the trial before.

Each participant had to switch their attentional focus between the same two streams and locations. The stream at the cued location was defined as target, the stream cued in the previous trial was defined as distractor. Crucially, this left one, never task-relevant stream and location for each participant, here defined as neutral. Between participants, we implemented three condition-to-location assignments to avoid any confound with the position of the neutral stream (neutral front (0°), neutral left (−45°) and neutral right (45°). We aggregated across the three condition-to-location assignments to obtain our measures of interest, i.e. neutral tracking of target, neutral and distractor. As the position of the neutral stream, the different audio streams were almost balanced between the 19 participants (neutral front: n=7; neutral right n= 6; neutral left n= 6).

Participants had to detect short repeats in the target stream. Each trial contained 6 repeats, which were randomly partitioned in the three streams. Before data collection, participants were familiarized with the experiment. During instruction, it was emphasized to respond as fast and accurately as possible to a repeat in the target stream, but also to listen to the content of the target stream. To familiarize participants with the repeats, we presented them a single sentence with one repeat included. They had to give oral feedback if they were able to detect the repeat. Further, we presented them with 6 training trials corresponding to the main experiment but using different audio streams. The main experiment consisted of 180 trials divided in 4 blocks, resulting in a total duration of 60 min. After each block, participants were able to take a rest. The total number of repeats was 360 per stream across the experiment.

We asked participants 15 multiple choice questions (with four possible answers, each) about the content of each audio stream at the end of the experiment. To avoid participants attending to the to-be-ignored audio stream, we did not ask the questions after every block. The order of the questions and the possible answers were randomized between participants.

### Behavioural data analysis

We evaluated participants’ behavioural performance in two ways. We analysed the proportion of detected repeats and, as a control, the proportion of correctly answered content questions.

We analyzed the detection of repeats in terms of signal detection theory. Button presses to repeats in a time window (150-1500 ms) after repeat onset were considered in this analysis. A button press following a repeat in the target stream was assigned as hit. Button presses following repeats in the distractor stream and in the neutral stream were assigned as separate types of false alarms. To differentiate between false alarms to repeats in the neutral versus distractor stream, we calculated sensitivity (d’) between hit rate and false alarms to distractor repeats [d’_target vs distractor_ = z(hit rate) – z(false alarm rate distractor)] and hit rate and false alarms to neutral repeats [d’ _target vs neutral_ = z(hit rate) – z(false alarm rate neutral)]. For this signal-detection analysis of repeats, we excluded one participant who did not respond to any repeats in the distractor stream.

A challenge in creating multiple-choice comprehension questions is to provide multiple (here: four) response options that cannot be solved based on prior knowledge or the possibility of excluding some of the response options. Hence, participants’ actual guess rate might be considerably higher than the theoretical chance level of 25 %. Thus, in a pilot experiment, we presented the multiple-choice comprehension questions to N=9 different participants who had not listened to the audio streams at all. This resulted in a new, ‘empirical’ chance level of 40 % (± 3.9 % S.E.M). In the following, we tested the proportion of correctly answered questions in the main experiment against this empirical chance level.

### Data acquisition and pre-processing

EEG was recorded using a 24 electrodes EEG-cap (Easycap, Herrsching, Germany; Ag–AgCl electrodes placed according to the 10-20 International System) connected to a SMARTING amp (mBrainTrain, Belgrade, Serbia). This is a mobile EEG system, which transfers the signal via Bluetooth to a recording computer(Waschke et al., 2017; Wöstmann, Waschke, et al., 2019).EEG activity was recorded with the software Smarting Streamer (mBrainTrain, version: 3.4.2) at a sample rate of 500 Hz. During recording, electrode FCz served as online reference and impedances were kept below 20 kΩ. No data loss was reported during the sessions.

Offline, EEG preprocessing was done using MATLAB (Version 2018a Mathworks Inc., Natick, MA, United States), built-in functions, custom-written code, and the Fieldtrip-toolbox (Oostenveld et al., 2011). EEG-data were re-referenced to the average of the electrodes M1 and M2 (left and right mastoids) and high- and low-pass filtered between 1 and 100 Hz (two-pass Hamming window, FIR). An independent component analysis (ICA) was computed on each participants’ EEG data. M1 and M2 were removed before ICA. ICA components related to eye blinks, eye movement, muscle noise, channel noise and line noise were identified by visual inspection and removed. On average, 8.37 of 22 (SD = 3.13) components were rejected. Components not associated with artifacts were back projected to the data. Clean EEG data were further processed. Neural speech tracking is associated with frequencies up to 8 Hz (Luo & Poeppel, 2007). Hence, EEG data were low-pass filtered again at 10 Hz (two-pass Hamming window, FIR). Afterwards, EEG data were resampled to 125 Hz and segmented into epochs corresponding to the trial length of 20s.

### Extraction of the speech envelope

The temporal fluctuations of speech were quantified by computing the onset envelope of each audio stream (Fiedler et al., 2017). First, we computed an auditory spectrogram (128 sub-band envelopes logarithmically spaced between 90-4000 Hz) using the NSL toolbox (Chi et al., 2005). Second, the auditory spectrogram was summed up across frequencies resulting in a broadband temporal envelope. Third, the onset envelope was obtained by computing the first derivative of this envelope. Finally, the onset envelope was down sampled to match the target sampling rate of the EEG analysis (125 Hz).

### Temporal response functions (TRFs)

The deconvolution kernel or impulse response, which describes the linear mapping between an ongoing stimulus to an ongoing neural response, is called the temporal response function (TRF). We used a multiple linear regression approach to compute the TRF (Crosse et al., 2016). More precisely, we trained a forward model using the onset envelopes (Fiedler et al., 2019) of the target, distractor, and neutral speech to predict the recorded EEG response. In this framework, we analysed time lags between –100 and +500 ms between envelope changes and brain response.

To account for the EEG variance attributable to the detection and processing of the behaviourally relevant repeats and corresponding evoked brain responses, we also included all onsets of the repeats in the three streams and the button press in the model as nuisance regressors, represented by stick functions. The onsets of the repeats are independent of the speech envelope regressors by design, since these were almost randomly (constraint of SNR threshold) added into the speech streams.

To prevent ill-posed problems and overfitting, we used ridge regression to estimate the TRF(Crosse et al., 2016). Lambda (λ) is the ridge parameter for regularization. We estimated the optimal ridge parameter that optimized the mapping between stimulus and response by leave-one-out cross-validation for each participant. First, the stimuli are segmented in M-trials and different ridge values (λ = 2^0^, 2^1^, … 2^20^) are predefined. In this approach, a separate model for each λ is calculated. Second, the trials are mixed, and each time one trial is left out. This trial is used as a test set, while the M-1 trials are used as a training set. Then, the models are averaged over the trials and convolved with the data from the matching test set to predict the neural response. This is done for every predefined λ. Computing the MSE between the predicted estimate and the original data provides a validation metric that enables to select the λ with the lowest MSE. We used the ridge value with the lowest MSE (specific for each subject) for the TRF model that jointly contained the target, distractor, and neutral onset envelopes as regressors.

TRFs were estimated based on the trials in the experiment. Participants had to switch their attention trial-wise between two of the streams. Hence, the trials enable the assignment of target, distractor and neutral onset envelope. Trial refers to the time window in which the stimulus and the response is cut to estimate the TRF. To avoid any conflicts with the cue, the first second of each trial was cut off in the EEG signal and the envelope onsets. One model was trained on 180 trials incorporating multiple predictor variables: the onset envelope for target, distractor and neutral stream, and the stick functions for the repeats and button presses using. Resulting in a single TRF for each predictor variable that predicts a separable response component. Similar to the TRF approach, we estimated TRFs for the embedded repeats but we modelled repeats as a stick function based on the repeat onset. To differentiate TRFs to speech and repeats, we refer to the TRF to repeats as regression-ERPs (rERPs; (Ehinger & Dimigen, 2019).

### Neural tracking

Neural tracking quantifies how strongly a single stream is represented in the EEG signal. TRFs were used to predict the EEG response. Predicted EEG response and measured EEG response were correlated using Pearson correlation, resulting in the neural tracking (r). We predicted the EEG signal on single trials using the leave-one-out cross-validation approach (see above). The resulting r-values were averaged across trials and participants. We obtained the neural tracking accuracy over TRF time lags by using a sliding-time window of time lags (size: 48 ms, 6 samples) with an overlap of 24 ms (3 samples) for the prediction (Fiedler et al., 2019; Hausfeld et al., 2018; Kraus et al., 2021; O’Sullivan et al., 2015). For every window position the neural tracking was calculated resulting in a time resolved neural tracking. We used the term “stream tracking” that refers to neural tracking of the envelope onsets, and “repeat tracking” that refers to the neural tracking of the repeat onsets. To obtain the repeat tracking, we used the same pipeline as for the speech tracking procedure (see above) with exception that we estimated neural tracking based on the onsets of repeats (instead of the speech onset envelope), which we modelled as stick functions.

### Statistical analysis

A study (Fiedler et al., 2019) investigated attentional effects of neural tracking in a comparable continuous speech paradigm by recording EEG of N=18 participants. It can be assumed that similar effect sizes achieved will be present again in a replication of auditory attention effects with the same sample size. The present study is supposed to detect neural tracking effects with at least a medium to large effect sizes (Cohen’s d ≥ 0.7) with a power of 80 % (two-sided, within-subject tests, Alpha = 0.05) for N = 18 subjects.

To answer the main research question (outlined in Fig.1b), we used generalized mixed models (*jamovi* 1.6, R 4.0). This approach enables us to include and jointly model factors that potentially influence behaviour and the neural response. These included at least the factor condition-to-location assignment (neutral front/ left/ right) and subject as random intercept.

To determine statistically significant differences in behavioural sensitivity (outcome measure), we included target versus the distractor stream and the target versus the neutral stream as categorical predictors in the model. To determine statistically significant differences in neural tracking (outcome measure), we included target, neutral and distractor stream as categorical predictors in the model. In both models, we included the factor condition-to-location assignment as covariate and the random intercept (subject ID) into the model. In order to derive Bayes factors, Bayesian t-tests were calculated (JASP Team, 2022).

For quantifying the brain-behaviour relations, we used a generalized linear mixed-effects model (repeat detected or not; binomial distribution, with logit link function). The predicted outcome variable was the binary response to the detection of a single repeat in the target stream (Hit=1/Miss=0). We included the encoding accuracies for the target, neutral and distractor stream as continuous, z-scored, *fixed-effects* predictors in our model. We assigned repeat tracking (trial-based) to each repeat within a trial. To control again for potential confounding between stream tracking and repeats, we also included the repeat tracking similar to the stream tracking in our model. Beside the factors *condition-to-location assignment* and *subject* as random intercepts, we also included *number of repeats during total experiment* and *number of repeats within a trial*, as well as *trial number* as random intercept into the model.

### Statistical analysis on time series

We were interested in finding time points that potentially differ in the time-resolved neural tracking (target enhancement: neutral vs. target and active suppression: neutral vs distractor) across subjects. We used a two-level statistical analysis, more specifically a cluster permutation test implemented in Fieldtrip (Oostenveld et al., 2011). Data from 22 channels were used in this analysis. As a test statistic at the single-subject level, we used one sample t-tests to test the time-resolved neural tracking to the target, neutral and distractor as well as the neutral-target, neutral-distractor, target-distractor difference against zero. At the group level, clusters were defined by the resulting t-values and a threshold that set to t-values that correspond to p < 0.05 for at least three neighbouring electrodes. Each observed cluster is compared to 5000 clusters with a permutation distribution. The permutation distribution was generated by randomly assigning the time-resolved neural tracking data to conditions. The relative number of iterations in which the summed t-statistic of the observed cluster is exceeded is indicated by the cluster p-value.

## Availability of data

The data that support the findings of this study are available from the corresponding author, MO, upon reasonable request.

